# Development and application of genetic ancestry reconstruction methods to study diversity of patient-derived models in the NCI PDXNet Consortium

**DOI:** 10.1101/2022.10.24.513591

**Authors:** Brian J. Sanderson, Paul Lott, Katherine Chiu, Juanita Elizabeth Quino, April Pangia Vang, Michael W. Lloyd, Anuj Srivastava, PDXNet Consortium, Jeffrey H. Chuang, Luis G. Carvajal-Carmona

## Abstract

Personalized medicine holds great promise for improving cancer outcomes, yet there is a large inequity in the demographics of patients from whom genomic data and models, including patient derived xenografts (PDX), are developed and for whom treatments are optimized. In this study we develop a genetic ancestry pipeline for the Cancer Genomics Cloud, which we use to assess the diversity of models currently available in the National Cancer Institute (NCI) supported PDX Development and Trial Centers Research Network (PDXNet). We show that there is an over-representation of models derived from patients of European ancestry, which is consistent with other cancer model resources. We discuss these findings in the context of disparities in cancer incidence and outcomes among demographic groups in the US. For example, for the top cancer health disparities affecting African Americans and Latinos, there is a significant lack of ethnic/race appropriate models needed to advance pre-clinical research and personalized clinical treatment. For stomach and liver tumors, which represent disparities in these two minority populations, there are only three available models derived from patients from such backgrounds. Fortunately, ongoing NCI-funded efforts in minority focused PDXNet centers are actively addressing these gaps. We further discuss these results in the context of power analyses to highlight the immediate need for the development of models from minority populations to address cancer health equity in personalized medicine.

## Introduction

Advances in our understanding of the genetic basis of cancers has led to the proliferation of so-called personalized treatments that can improve outcomes^1^, and yet there are disparities in the groups for which these treatments are most impactful. A major contributor to these disparities is an over-representation of donors with European ancestry in cell lines, sequence data, and patient-derived models of human cancer.^2^ Health disparities in cancer incidence and outcome are pervasive in minority communities in the US, and are driven by complex interactions between socioeconomic factors and potentially also by genetic variants that differ in frequency between demographic groups^3–5^. Understanding whether and how germline and somatic variants influence cancer health disparities will require a greater investment in the development of cancer models that better represent the diversity of human genetic and epigenetic variation.

Genetic ancestry, which describes the relationships between individuals and populations based on shared genetic history, can also provide insight into variance in disease risk, prognosis, and response to therapies. For example, studies in Latinos, who trace their ancestry to Spanish conquistadors, African slaves and Indigenous American women, have shown that Indigenous American ancestry has an inverse relationship with breast cancer risk^6,7^. This is consistent with epidemiological data suggesting a low incidence of breast cancer in Latin American countries with large numbers of Indigenous Americans^8^. Interestingly, we showed recently that Indigenous American ancestry is associated with tumor phenotypes such as *ERBB2* amplification^9^, while other studies have shown similar associations between indigenous ancestry and EGFR-mutated lung tumors^10^. Studies in African American (AA) populations have also shown that African ancestry is associated with higher risk of prostate cancer^11^, and that genetic variants exclusively found in people of African ancestry are associated with the prevalence of triple negative breast tumors, which are more aggressive and common in AA patients^12^. Despite the important differences between demographic groups revealed in the above epidemiological studies, a major limitation in the field has been the paucity of genetic germline and somatic data from patients belonging to diverse populations^13^. Furthermore, as discussed in our recent commentary, another major limitation is the lack of models derived from minority patients because such models recapitulate their genome and epigenome and are needed to advance precision health equity in such populations^2^. To address these research gaps, the National Cancer Institute (NCI) has supported two centers within the PDX (Patient-Derived Xenograft) Development and Trial Centers Research Network (PDXNet) to work with minority populations to create models that reflect their genetic and environmental history, to characterize their genome, and to facilitate studies that will lead to future minority-focused clinical trials with the goal of addressing current cancer health disparities.

Several algorithms exist to estimate ancestry from genetic data, which can be roughly categorized as model-based methods that rely on statistical inference based on, for example, admixture linkage disequilibrium, and distance-based methods that estimate ancestry using clustering, network theory, or graph theory.^14,15^ STRUCTURE was a trailblazing program that led to a proliferation of parametric methods that model and partition linkage disequilibrium due to admixture to ascertain the genetic structure of populations, or to identify the proportional contribution of different reference populations to the genetic makeup of individuals.^16,17^ Although this type of model-based method can provide robust estimates of ancestry, they often require detailed population or pedigree information that may not be readily available from protected or anonymized patient data, and they also tend to have long run-times and detailed hands-on analyses that make them less tractable for pipeline development.^18^ Alternatively, distance-based approaches use methods such as principal component analysis (PCA)^19,20^ to assign genetic ancestry by projecting samples into high-dimensional space and comparing their positions relative to data from reference populations, which is a much less computationally intensive task than most model-based methods.

In this manuscript, we describe the development of an ancestry estimation pipeline on the Seven Bridges Genomics (SBG) Cancer Genomics Cloud (CGC) as part of the NCI-PDXNet project. This pipeline implements the program SNPweights,^21^ which uses a distance-based method to estimate genetic ancestry by weighting factors for annotated SNPs derived from a PCA of genetic data from reference populations. Using reference weights to assign proportions of genetic ancestry requires much less computational time than running the initial PCA on the reference data to derive the weights. This allowed us to implement this program alongside the standard sample intake pipeline for whole exome data. We use this new pipeline to describe the diversity of samples currently in PDXNet in terms of the genetic ancestry backgrounds of patients from whom the tumors were sampled. We further present power analyses to argue that additional PDX models are critically needed to address the disparities in cancer treatment outcomes in patients from minority backgrounds in the United States.

## Methods

### SNPweights Panel Design

SNPweights v 2.1 includes several weight panels that differ in the representation of continental reference populations.^8^ However, in our benchmark testing of published SNPweights models, we found that the “NA” panel, which includes reference data from African, East Asian, European, and Indigenous American populations, yielded ancestry estimates that were not concordant with ADMIXTURE estimates using 1000 Genomes and Indigenous American references. We designed a new reference panel for the program SNPweights to improve the classification of individuals from ancestral backgrounds that originated in the Americas and South Asia. We used as a reference set human genetic data that were downloaded from the 1000 Genomes Project Phase III,^22^ GenomeAsia 100K^23^, and INMEGEN^24^, with data imputed using the TOPmed Imputation Server.^25^ We estimated continental genetic ancestral fractions for each sample as fractions of five continental ancestral categories, which include European (EUR), African (AFR), Indigenous American (AMR), East Asia (EAS), and South Asian (SAS)^26^.

Reference data were converted from the Genome Reference Consortium (GRC) GRCh37 build of the human genome^27^ to the GRCh38 build^28^ using UCSC LiftOver.^29,30^ The raw data consisted of 4,471,291 SNPs identified in 11,448 individuals (Table S1). A principal component analysis was performed on the filtered data using SMARTPCA in the EIGENSOFT package v. 7.2.0^19,20^ to identify 1,990 individuals with little to no admixture, which formed tight clusters at the termini of the ancestral axes (238 African, 348 Indigenous American, 408 European, 766 East Asian, 230 South Asian; Fig. S1). Markers were filtered for minor allele frequency ≤ 5%, Hardy-Weinberg equilibrium P ≤ 0.0005, independent pairwise linkage disequilibrium pruning using a 50Kbp window a step size of 5 markers and r^2^ threshold of ≥ 0.5, and exclusion of high LD regions, leaving 264,153 SNPs.

The filtered data were converted into EIGENSTRAT format using convertf in the EIGENSOFT package v. 7.2.0.^19,20^ A principal component analysis was performed on the filtered data using SMARTPCA in the EIGENSOFT package v. 7.2.0. Finally, the weighting factors for ancestry inference were extracted from the PCA results using the program calc_snpwt.py from the SNPweights package v 2.1.

This model was validated and benchmarked using 2,387 admixed individuals with ADMIXTURE v1.3.0^31^ in both supervised and unsupervised modes. In our assessment of the original SNPweights NA panel’s estimates of Asian ancestry, individuals identified by 1000 Genomes as SAS differed on average by 0.5507 (SD:0.0910) in comparison to ADMIXTURE supervised. Among the same 1000 Genomes individuals identified as AMR and SAS, the new SNPweights ancestral estimates differed from ADMIXTURE supervised by mean of 0.0231 (SD: 0.0167) and 0.0369 (SD: 0.0400), respectively.

### Ancestry Estimation

To assess the diversity of genetic backgrounds represented by the PDX models in PDXNet, we estimated genetic ancestry using the Binary Alignment Map (BAM) files that were prepared as part of the standard intake process for whole exome data on the CGC.^32^ Genotypes were estimated using the programs bcftools mpileup with the parameters -I -q 30 -Q 20 and bcftools call with the parameter -m in the bcftools package v. 1.9.^33^ These genotypes were then filtered using bcftools filter with the parameters -g3 -G10 -e’%QUAL<20 || (RPB<0.1 && %QUAL<25) || (AC<2 && %QUAL<25) || INFO/DP>70 || INFO/DP<4’. The filtered genotypes were annotated with SNP IDs from dbSNP build 151^19^ using bcftools annotate. The annotated genotypes were converted into EIGENSTRAT format using the program vcf2eigenstrat.py in the Genetic Data Conversion package.^35^ Finally, genetic ancestry proportions were estimated using the program inferanc in the SNPweights package v 2.1. Samples were then categorized into the ancestry group for which the proportion was greater than 0.7. When no category scored greater than 0.7 the samples were labeled MIXED, and were categorized based on the top two ancestry groups for those samples.

### Power Analyses

To determine the power to detect driver mutations that differ in frequency between demographic groups, we performed a power analysis using the Power for Genetic Association Analyses (PGA) package.^36^ Statistical power was modeled for a dominant mode of inheritance, assuming that markers and causative SNPs are in complete linkage (LD = 1), and that there were 200 effective degrees of freedom, which would account for 200 comparisons due to multiple genetic markers. The causative mutation was modeled to be absent in one group, and present at varying low frequencies of 0.075, 0.1, 0.15, 0.2, 0.25.

We were also interested to determine the sampling effort necessary to develop new PDX models for a newly discovered driver mutation that is segregating at a known frequency in a minority population. We modeled the sampling effort necessary to identify at least five individuals with the driver mutation using the hypergeometric distribution across varying frequencies from 0.01 to 0.25 using the function phyper in R.^37^ Plots from both power analyses were prepared using the package ggplot2 in R.^38^

### Cancer Health Disparities Data Analyses

We used data from the NCI Surveillance, Epidemiology, and End Results program from 18 population-based cancer registries (SEER18) from 2013-2017 to identify the top 10 causes of cancer mortality in AA and Latinos, the two largest US minority populations, and in Non-Latino Whites (NLW). As PDXs are primarily used for pre-clinical studies to develop new therapies, we focused on analyses in cancer mortality disparities. In these analyses, once the top ten causes of cancer mortality in men and women from each of the three racial/ethnic groups were identified, we estimated the fold difference in mortality between minority populations and NLW, which we termed the disparity ratio. Cancers for which mortality rates ranked exclusively high in minorities or that have disparity ratios > 1 were then designated as “priority malignancies” for model development. We then queried the PDXNet dataset to identify the number of ethnic/race appropriate models for such priority malignancies and estimated the number of models available.

## Results

### Genetic Ancestry Among PDX Models in PDXNet

The first three principal components explained 44.63%, 24.71%, and 17.6% of the variation of the filtered non-admixed ancestral reference genotype matrix, respectively. Samples cluster well by continental ancestry category among these three principal components (Fig. S1), with the exception of the AMR and SAS categories which show less tight clustering, potentially due to population diversity or a lack of genome-wide references with regards to AMR.

The models available in PDXNet as of September 2022 represented 980 unique patients, and self-reported race and ethnicity (SIRE) information was available for 595 patients. These models were developed by the six PDX Development and Testing Centers in the PDXNet Consortium, as well as models from the NCI Patient-Derived Models Repository (PDMR). Our genetic ancestry pipeline estimated 65 models with majority African ancestry and 1 model with majority American ancestry (Fig. 1A), as well as 13 models with mixed African and American and 42 models with mixed American and European ancestry (Fig. 1B). The estimates of genetic ancestry proportions for these samples were highly concordant with SIRE information (Table S2).

**Figure 1.**
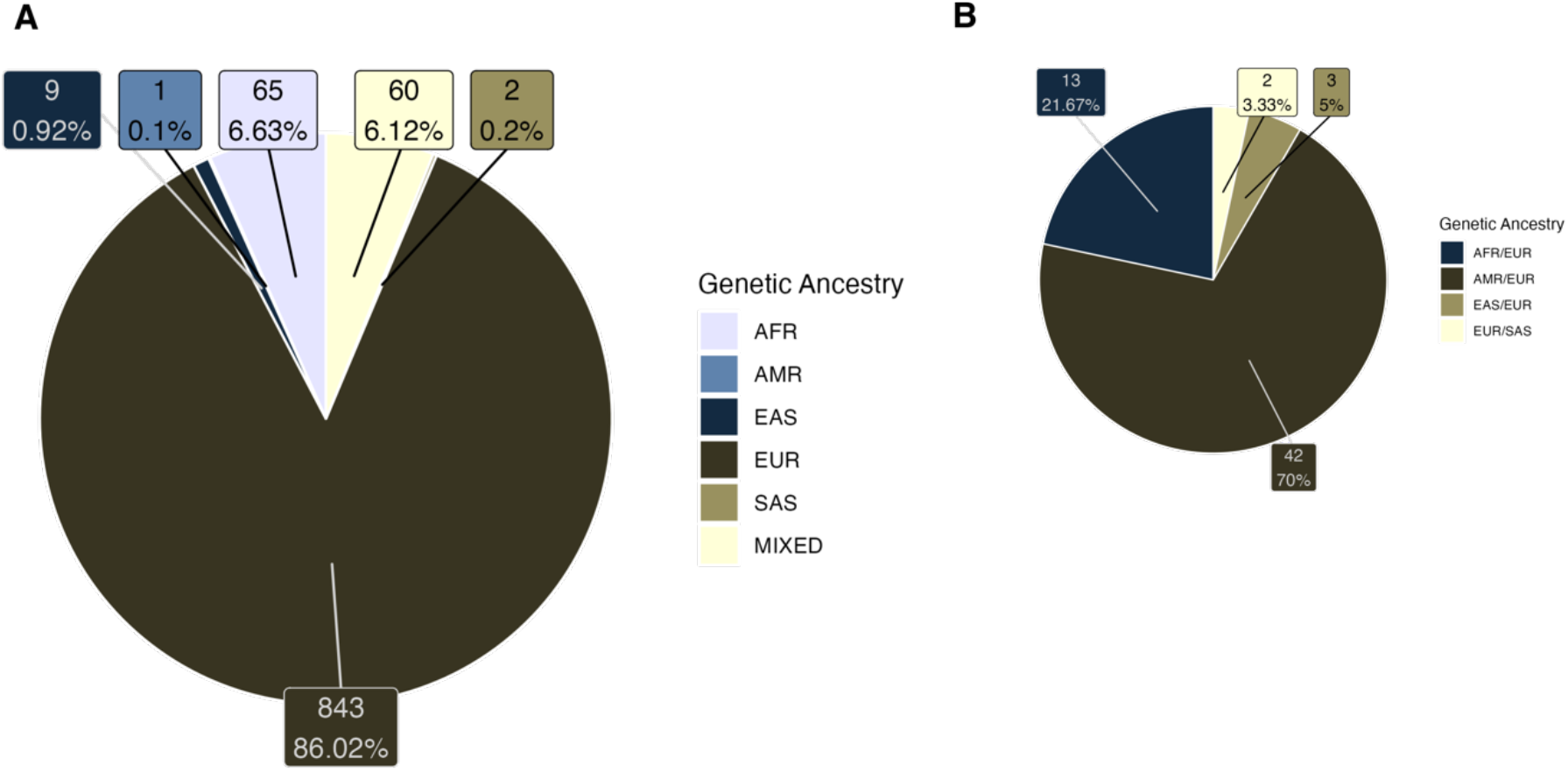
Diversity of genetic ancestry estimates from PDXNet models. A: Inferred genetic ancestry for 980 models across all cancer types. B: Top two categories for 60 “MIXED” samples with no category > 70%. Numbers and percentages in each box reflect the models assigned to that category based on estimates from the SNPweights analysis.

Overall, a majority of the models in PDXNet originate from patients with majority European continental ancestry (Fig. 1). Certain cancer types, such as breast cancers (Fig. S2), have a greater representation of patients from non-European backgrounds due to the efforts of minority PDX testing and development centers at the Baylor College of Medicine and University of California Minority Patient-Derived Xenograft Development and Trial Center, which are members of the PDXNet consortium.

### Power to Detect Drivers and Develop New PDX Models

Our power analysis demonstrates that a study with 150-300 each in of two ancestry categories would have > 80% to detect the presence of a dominant driver mutation that segregates between them (Fig. S3). We also found that for a known driver mutation that segregates at frequencies of 0.01 – 0.1 in a given population, sampling 50-100 individuals from that population should be sufficient to identify at least five patients from which PDX models could be developed (Fig. S4).

### Implications for cancer health disparities

In a recent commentary, we argued for the need to diversify patient-derived models and the implications of the limited diversity of such models to advance precision medicine in minority populations.^2^ To contextualize the proportional representation of models from these groups in PDXNet, we obtained cancer mortality and disparities data for African Americans and Latino reported to SEER18^39,40^. Because race and ethnicity are complex and not reducible to proportions of genetic ancestry, we compared the cancer disparities data with self-reported race and ethnicity from the model donors for which those data are available. In Table 1, we show data from the ten most common causes of female cancer mortality in NLW, AA and in Latinos, ranked by the incidence in NLW. In Table 2, we show similar data for males. While some of the leading causes of cancer mortality are common but have different rankings for NLW, AA and Latinos, others are particularly high in minorities and usually associated with disparities. For example, common female causes of cancer-related death such as those affecting the lung and bronchus, breast, colon and rectum, pancreas, ovary, corpus uterus, leukemias and liver and intrahepatic bile duct affect all American women. American men, on the other hand, experience high mortality rates for tumors in the lung and bronchus, prostate, colorectum, pancreas, liver and intrahepatic bile duct, leukemias, urinary bladder and non-Hodgkin lymphoma. These tumors rank in the top ten for all three racial/ethnic groups. However, there are several malignancies with a disproportionally high burden in minority populations or that are less common in NLW. Female mortality data show a disproportionate stomach cancer burden for Latinos and AA and multiple myeloma for AA (Table 1). AA men disproportionally died due to stomach tumors and multiple myeloma while Latino men are particularly affected by stomach and renal tumors (Table 2). For other common tumors, these two minority groups also show important cancer health disparities when compared to NLW (see disparity ratio columns in Tables 1 and 2). These cancer mortality disparities include liver and intrahepatic tumors for Latinos (with mortality rates > 47% when compared to NLW) and AA (with mortality rates > 23% when compared to NLW) in both sexes; colorectal and pancreatic tumors for both sexes in AA (with mortality rates > 28% and > 18%, respectively, when compared to NLW); lung and prostate tumors for AA men (with mortality rates 16% and 208%, respectively, higher than in NLW); and breast and uterine tumors for AA women (with mortality rates 40% and > 91%, respectively, higher than in NLW).

**Table 1.** Top 10 Female Age-Adjusted Mortality Rates. SEER18 2013-2017

| Site | Non-Hispanic Whites |  |  | African Americans |  |  |  | Latin Americans |  |  |  |
| --- | --- | --- | --- | --- | --- | --- | --- | --- | --- | --- | --- |
|  | Rank | AA-MR | Count | Rank | AA-MR | Count | DR | Rank | AA-MR | Count | DR |
| Breast | 1 | 44.35 | 84,996 | 1 | 53.18 | 14,211 | **1.20** | 1 | 28.58 | 9,925 | 0.64 |
| Lung and Bronchus | 2 | 38.73 | 71,424 | 2 | 36 | 9,726 | 0.93 | 2 | 16.83 | 5,451 | 0.43 |
| Colon and Rectum | 3 | 19.92 | 39,257 | 3 | 25.82 | 6,793 | **1.30** | 3 | 14.77 | 4,932 | 0.74 |
| Corpus and Uterus | 4 | 9.39 | 17,893 | 4 | 13.34 | 3,657 | **1.42** | 5 | 7.6 | 2,652 | 0.81 |
| Pancreas | 5 | 8.9 | 16,697 | 5 | 11.87 | 3,200 | **1.33** | 4 | 8.46 | 2,811 | 0.95 |
| Lymphoma | 6 | 8.34 | 16,126 | 7 | 6.63 | 1,735 | 0.79 | 6 | 7.5 | 2,426 | 0.90 |
| Ovary | 7 | 7.47 | 13,569 | 8 | 6.27 | 1,730 | 0.84 | 7 | 5.99 | 2,158 | 0.80 |
| Leukemia | 8 | 5.81 | 10,884 | 10 | 5.39 | 1,409 | 0.93 | 10 | 4.49 | 1,636 | 0.77 |
| Skin excluding Basal and Squamous | 9 | 5.55 | 10,881 | 28 | 0.63 | 162 | 0.11 | 20 | 1.61 | 535 | 0.29 |
| Urinary Bladder | 10 | 5.16 | 10,443 | 13 | 4.31 | 1,091 | 0.84 | 15 | 2.54 | 781 | 0.49 |
| Myeloma | 14 | 2.89 | 5,448 | 6 | 7.68 | 2,002 | **2.66** | 13 | 3.16 | 1,019 | **1.09** |
| Kidney and Renal Pelvis | 11 | 4.52 | 8,619 | 9 | 5.48 | 1,426 | **1.21** | 11 | 4.41 | 1,462 | 0.98 |
| Cervix Uteri | 16 | 2.73 | 4,208 | 11 | 5.18 | 1,427 | **1.90** | 12 | 3.84 | 1,497 | **1.41** |
| Liver and Intrahepatic Bile Duct | 15 | 2.86 | 5,265 | 14 | 4.23 | 1,217 | **1.48** | 8 | 5.97 | 2,004 | **2.09** |
| Stomach | 17 | 2.29 | 4,284 | 12 | 5.04 | 1,317 | **2.20** | 9 | 5.52 | 1,959 | **2.41** |

Abbreviations: AA-MR - Age Adjusted Mortality Rate; DR - Disparity Ratio. Count is 5-year number of cases. Database: Incidence-Based Mortality - SEER Research Data, 18 Registries, Nov 2019 Sub (2000-2017).

**Table 2.** Top 10 Male Age-Adjusted Mortality Rates. SEER18 2013-2017.

| Site | Non-Hispanic Whites |  |  | African Americans |  |  |  | Latin Americans |  |  |  |
| --- | --- | --- | --- | --- | --- | --- | --- | --- | --- | --- | --- |
|  | Rank | AA-MR | Count | Rank | AA-MR | Count | DR | Rank | AA-MR | Count | DR |
| Prostate | 1 | 66.63 | 92,170 | 1 | 125.6 | 19,303 | **1.89** | 1 | 51.2 | 10,477 | 0.77 |
| Lung and Bronchus | 2 | 49.97 | 75,245 | 2 | 60.7 | 11,920 | **1.21** | 2 | 25.15 | 6,157 | 0.50 |
| Colon and Rectum | 3 | 28.92 | 41,546 | 3 | 37.88 | 6,935 | **1.31** | 3 | 23.56 | 6,060 | 0.81 |
| Urinary Bladder | 4 | 24.09 | 33,566 | 6 | 12.68 | 1,990 | 0.53 | 6 | 10.37 | 2,189 | 0.43 |
| Skin excluding Basal and Squamous | 5 | 15.79 | 22,136 | 21 | 1.24 | 212 | 0.08 | 16 | 2.52 | 629 | 0.16 |
| Lymphoma | 6 | 14.09 | 20,030 | 8 | 10.92 | 2,181 | 0.78 | 5 | 11.55 | 3,041 | 0.82 |
| Pancreas | 7 | 11.46 | 17,479 | 5 | 13.78 | 2,840 | **1.20** | 7 | 9.6 | 2,669 | 0.84 |
| Leukemia | 8 | 10.91 | 15,398 | 11 | 8.24 | 1,529 | 0.76 | 10 | 7.13 | 2,143 | 0.65 |
| Kidney and Renal Pelvis | 9 | 10.39 | 15,294 | 7 | 12.31 | 2,321 | **1.18** | 8 | 9.53 | 2,566 | 0.92 |
| Liver and Intrahepatic Bile Duct | 10 | 8.35 | 13,623 | 4 | 13.88 | 3,375 | **1.66** | 4 | 15.79 | 4,955 | **1.89** |
| Myeloma | 14 | 5.14 | 7,496 | 9 | 10.54 | 1,903 | **2.05** | 11 | 4.96 | 1,239 | 0.96 |
| Stomach | 13 | 5.61 | 8,325 | 10 | 10.29 | 1,956 | **1.83** | 9 | 9.42 | 2,624 | **1.68** |

Abbreviations: AA-MR - Age Adjusted Mortality Rate; DR - Disparity Ratio. Count is 5-year number of cases. Database: Incidence-Based Mortality - SEER Research Data, 18 Registries, Nov 2019 Sub (2000-2017).

In Table 3, we show the number of PDX models available in PDXNet derived from NLW, AA and Latino patients (based on self-reported race and ethnicity) with the “priority malignancies” identified above (liver, stomach, kidney/renal pelvis, colon and rectum, pancreas, lung, prostate, breast, corpus and uterus and multiple myeloma). For stomach tumors, a malignancy that disproportionally affects both minority groups, only one appropriate PDX exists for AA and another one for Latinos. For liver cancer, another malignancy with high burden in both groups, there are no race/ethnic appropriate models for AA and only one for Latinos in PDXNet. Kidney tumor models from Latinos also lacks race/ethnic appropriate models. The number of models for malignancies with high burden in AA is also dismal with no race appropriate models for multiple myeloma or prostate cancer, four for pancreas cancer, three for endometrial cancer, four for colorectal cancer and two for lung cancer. Only breast cancer is moderately represented for AAs, with 14 models from this minority group currently available in PDXNet. While cancers that are rare in NLW, such as multiple myeloma and stomach, liver and kidney tumors, also have low numbers for this majority racial group, the disparities for common cancers in NLW are striking. For example, there is only one colorectal cancer PDX from Latinos for every ~36 from NLW. For each African American colorectal cancer PDX, there are 27 from NLW. NLW pancreas and lung cancer PDXs are also 7-fold and 13-fold higher than for AA, respectively. These analyses therefore not only identify disparities in existing models but also highlight that developing more diverse pre-clinical models is needed to address cancer mortality disparities. Given the disproportionally high burden caused by liver and stomach tumors in AA and Latinos, by renal tumors in Latinos, by colorectal, pancreas, lung, prostate, breast and uterine tumors and multiple myeloma in AA, we suggest that they should represent a priority for model development in PDXNet and similar initiatives.

**Table 3.**
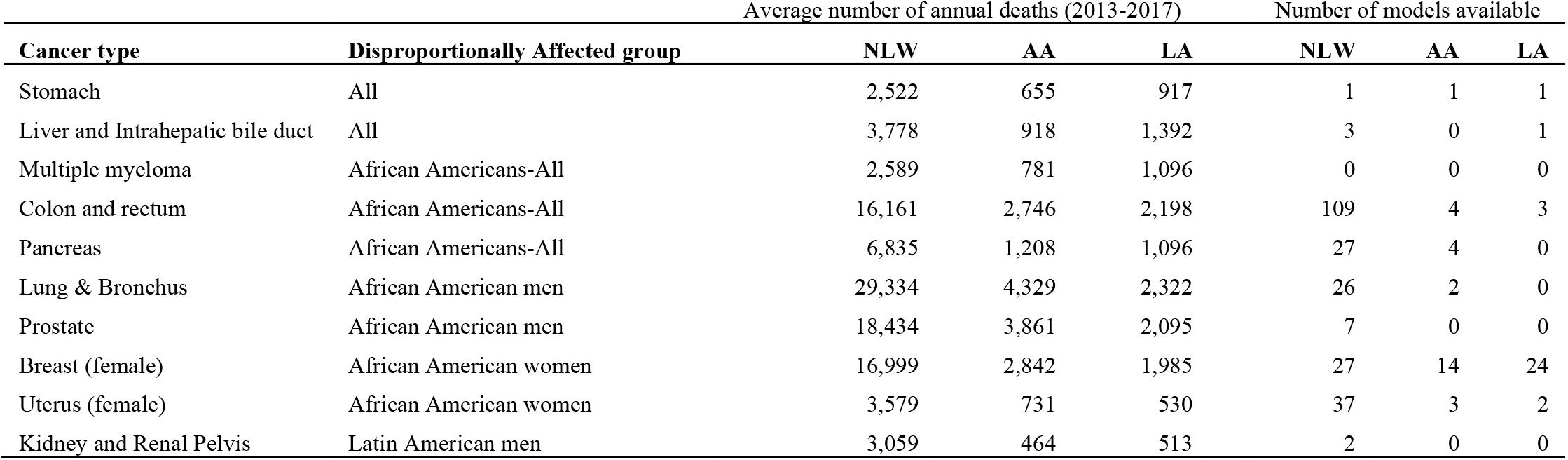
Summary of annual deaths and PDX model availability

## Discussion

In this study we developed a genetic ancestry pipeline for the Cancer Genomics Cloud, which we used to assess the diversity of models currently in PDXNet. We found that our genetic ancestry estimates had a high correspondence with self-reported race and ethnicity, and thus are useful for comparing the diversity of models to published cancer health disparities. Patients with European genetic ancestry are highly overrepresented in the PDXNet models, which reflects similar disparities in other cancer resources^2^. Recent efforts of the PDXNet have yielded an increase in 18 models of breast cancers likely derived from patients with African genetic ancestry, and 26 models likely derived from Latin American genetic ancestry, which represent a 4-fold and 26-fold increase in models above those previously available, respectively. Furthermore, with the addition of two minority and disparity focused centers in PDXNet in late 2018, there several models from minority patients that will be soon deposited in the PDMR, including around 20 new gastric cancer models from self-reported Latinos. Yet, as our power analyses showed, there are still far fewer models than would be required to conduct a study with sufficient power to identify genetic variants with biologically relevant effects on the cancer health disparities between demographic groups. This highlights the critical need for further investment in model development to help reach health equity goals.

Health disparities between demographic groups in the US are complex, and involve the interaction of many factors including structural inequity, socioeconomic status, and variance in exposures to environmental harms.^3–5^ Our focus on genetics in this study is motivated by our belief that personalized medicine holds great promise for improving patient outcomes, and that this promise should be realized equitably. The evolution of cancers leverages the germline and somatic background of patients in which tumorigenesis occurs, and the progression of cancer depends on the interactions between somatic mutations and the normal tissue in the microenvironment on which it grows.^41^ It is therefore important to understand the diversity of genetic backgrounds in which cancers evolve to identify relevant biomarkers that could help to address treatment disparities. We want to emphasize that human genetic diversity is complex, there is large variance within any grouping of human populations, and we do not seek to naively assign risk factors for cancer incidence to broad, biologically dubious categories.^42–44^ Rather, we hope this work will help to motivate and facilitate an understanding of whether personalized medicine approaches have the potential to help address current cancer health disparities by increasing the number and diversity of models available to identify whether and how segregating germline and somatic variants can help explain those disparities.

Categorization of models based on continental ancestry has several limitations worth discussing. The concept of continental ancestry is premised on the concept of continental races, the biological and biomedical relevance of which are debated and controversial.^42,45^ The history of human migration and gene flow over the past 100,000 years is complex and cannot be circumscribed by continental borders.^43,44^ Additionally, analyses based on categorization of individuals by SIRE or ancestry likely elide the complex interactions of demography, the environment, and socioeconomic factors^3,4,44^ Our goal in this study was to help motivate the critical need for cancer models that better reflect the diversity of human genetic variation in order to help achieve equity in personalized medicine. In this way, we seek to better understand the extent to which genetic variants that segregate among demographic groups impact cancer health disparities.

In conclusion, we developed and implemented a pipeline for genetic ancestry evaluations in publicly available patient derived models and highlight the fact that most of them have a predominantly European genetic ancestry. We estimated that to understand biological differences and responses to therapy between different genetic ancestries, hundreds of models are needed, highlighting the need to diversify the models. Furthermore, using self-reported race/ethnic information available in ~50% of the models and cancer health disparities data in the two largest US minority populations, we showed that there very few or, in many cases, no models to develop therapies for the cancer types with the highest burden in minority patients. While ongoing efforts promise to diversify these available models, we encourage funders to support additional efforts aimed at developing and characterizing new models that equitable help realize the promise of cancer precision medicine to all Americans.

## Acknowledgements

This work was primarily funded by grants from the Cancer Moonshots program (NCI U24-CA224067, NCI U54-CA224083, NCI U54-CA224070, NCI U54-CA224065, NCI U54-CA224076, NCI U54-CA233223, NCI U54-CA233306). LGC-C is grateful for funding received from The Auburn Community Endowed Chair in Basic Cancer Research; The California Initiative to Advance Precision Medicine (contract OPR18111), The Heart, BrEast, and BrAin HeaLth Equity Research (HEAL HER) program and from the National Cancer Institute (grants R01CA223978, R21CA199631, and P30CA093373) and National Center for Advancing Translational Science (UL1TR001860) of the National Institutes of Health. JEQ and APV received diversity supplement support from the National Cancer Institute (R01CA223978-2S2R01and R01CA223978-05S2). The content is solely the responsibility of the authors and does not necessarily represent the official views of the National Institutes of Health.

## Data Accessibility

The new SNPweights panel will be made publicly available upon acceptance of this manuscript. The ancestry estimation pipeline is implemented in the Common Workflow Language and will be made available in the public applications gallery of the Cancer Genomics Cloud upon acceptance of this manuscript.

## Supplementary Figure Legends

**Figure S1.**
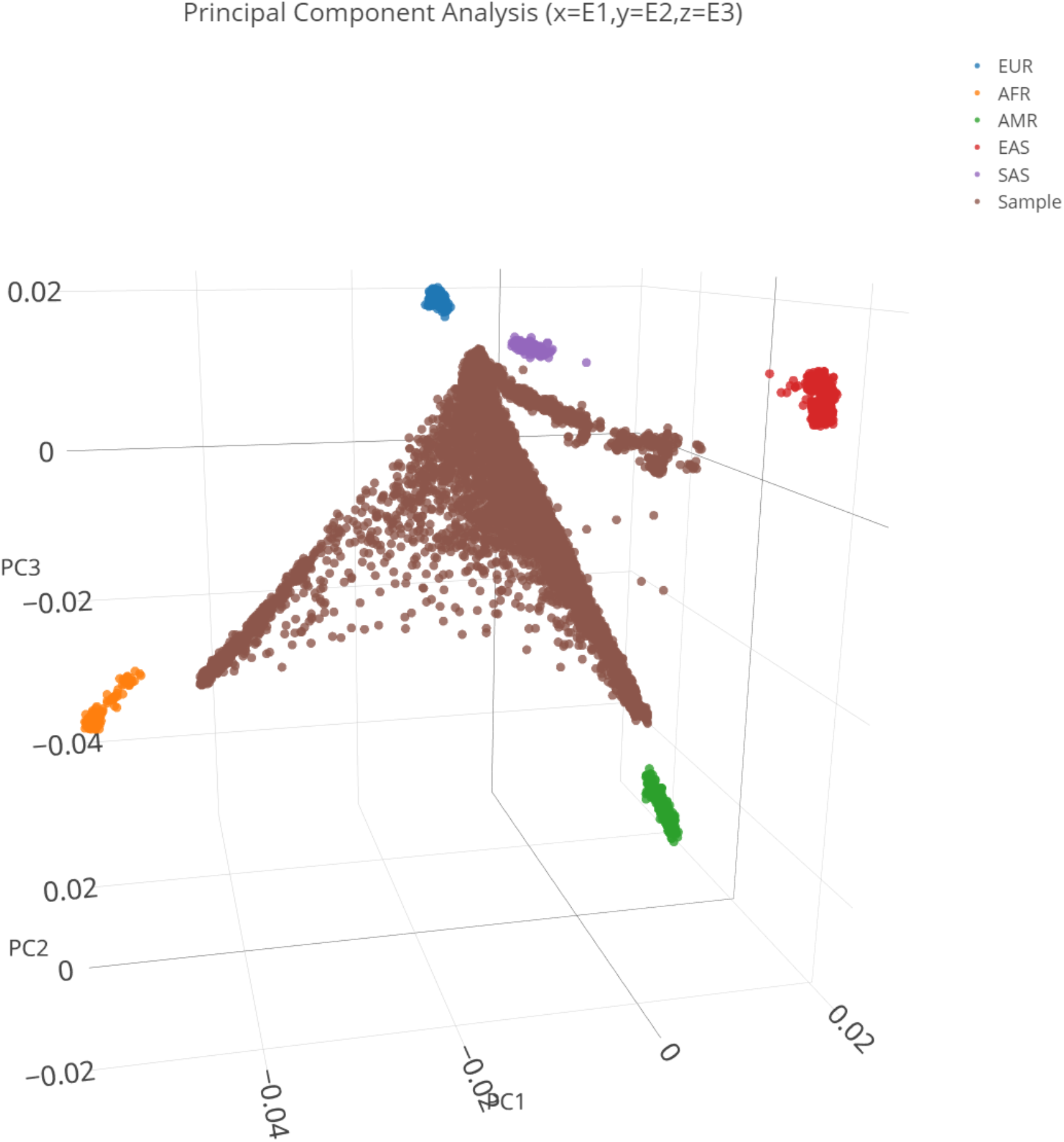
Clustering of 1,990 non-admixed reference samples and 2,387 admixed samples (brown) individuals based on the first three principal components. Colors represent the continental ancestry category assigned to each sample based on the country of origin.

**Figure S2.**
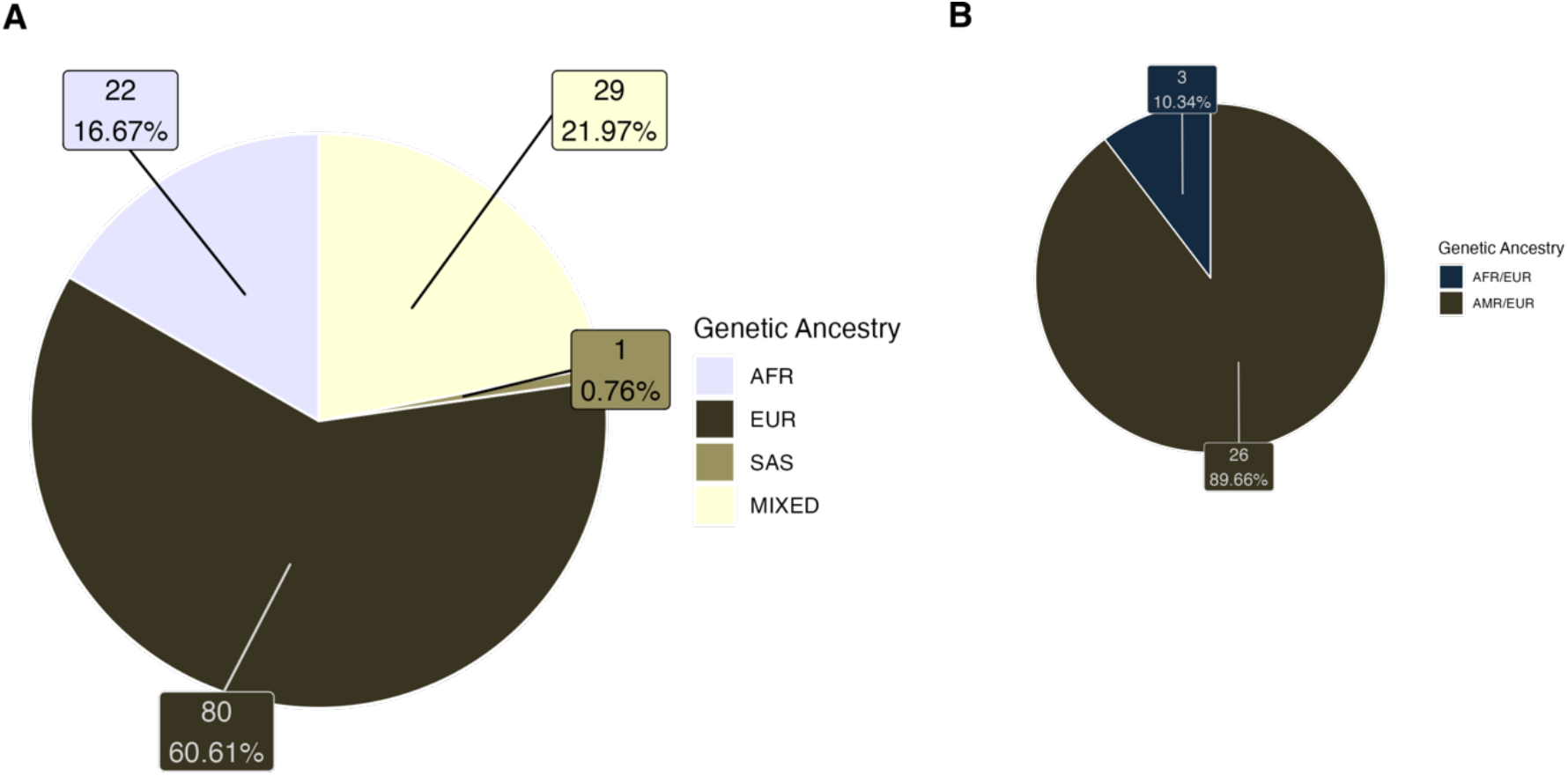
Diversity of genetic ancestry estimates from PDXNet breast cancer models. A: Inferred genetic ancestry for 132 breast cancer models in PDXnet. B: Top two categories for 29 “MIXED” samples with no category > 70%. Numbers and percentages in each box reflect the models assigned to that category based on estimates from the SNPweights analysis.

**Figure S3.**
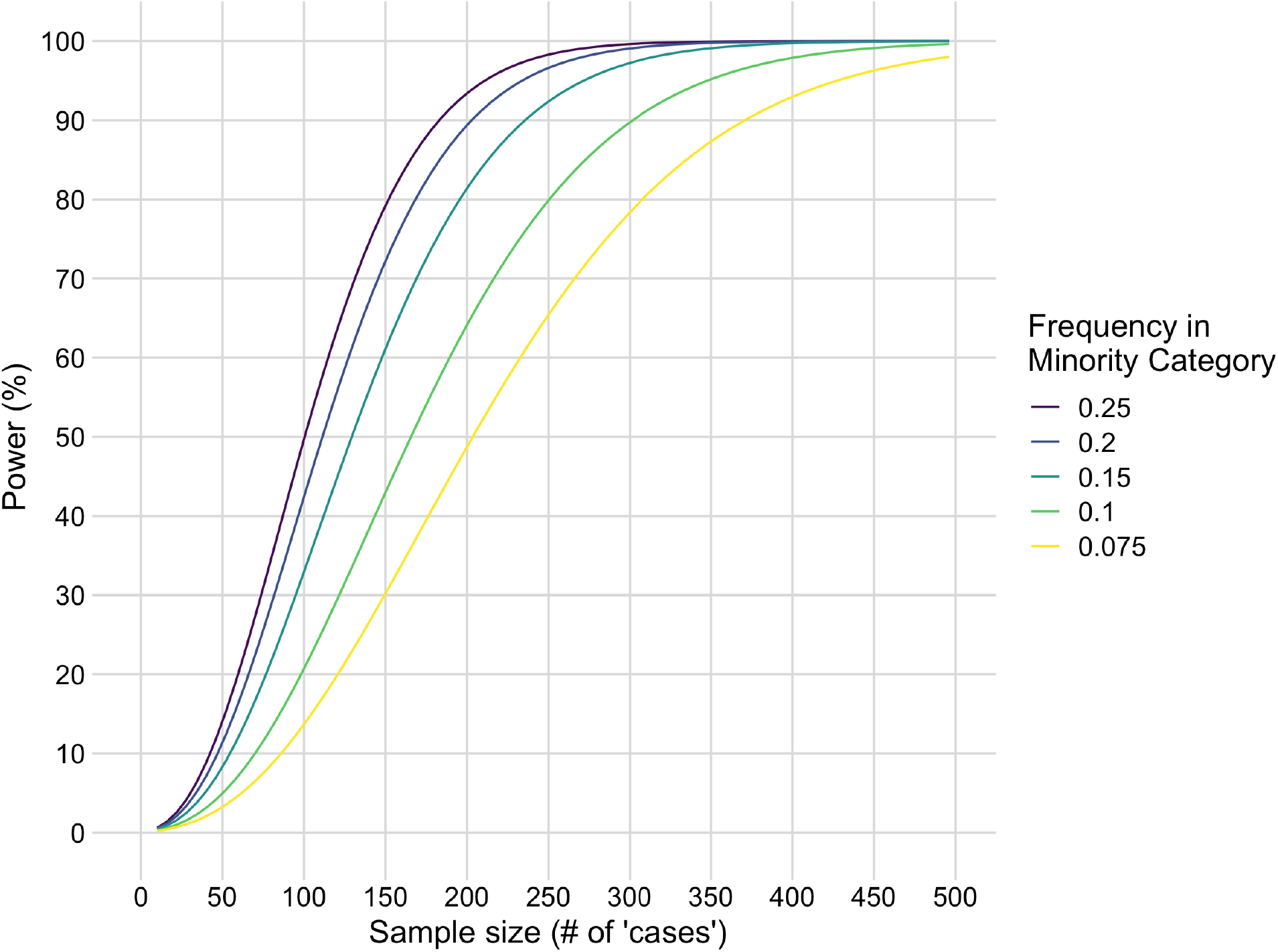
Power to detect a driver mutation that is absent in EUR but present in a non-EUR category at varying low frequencies.

**Figure S4.**
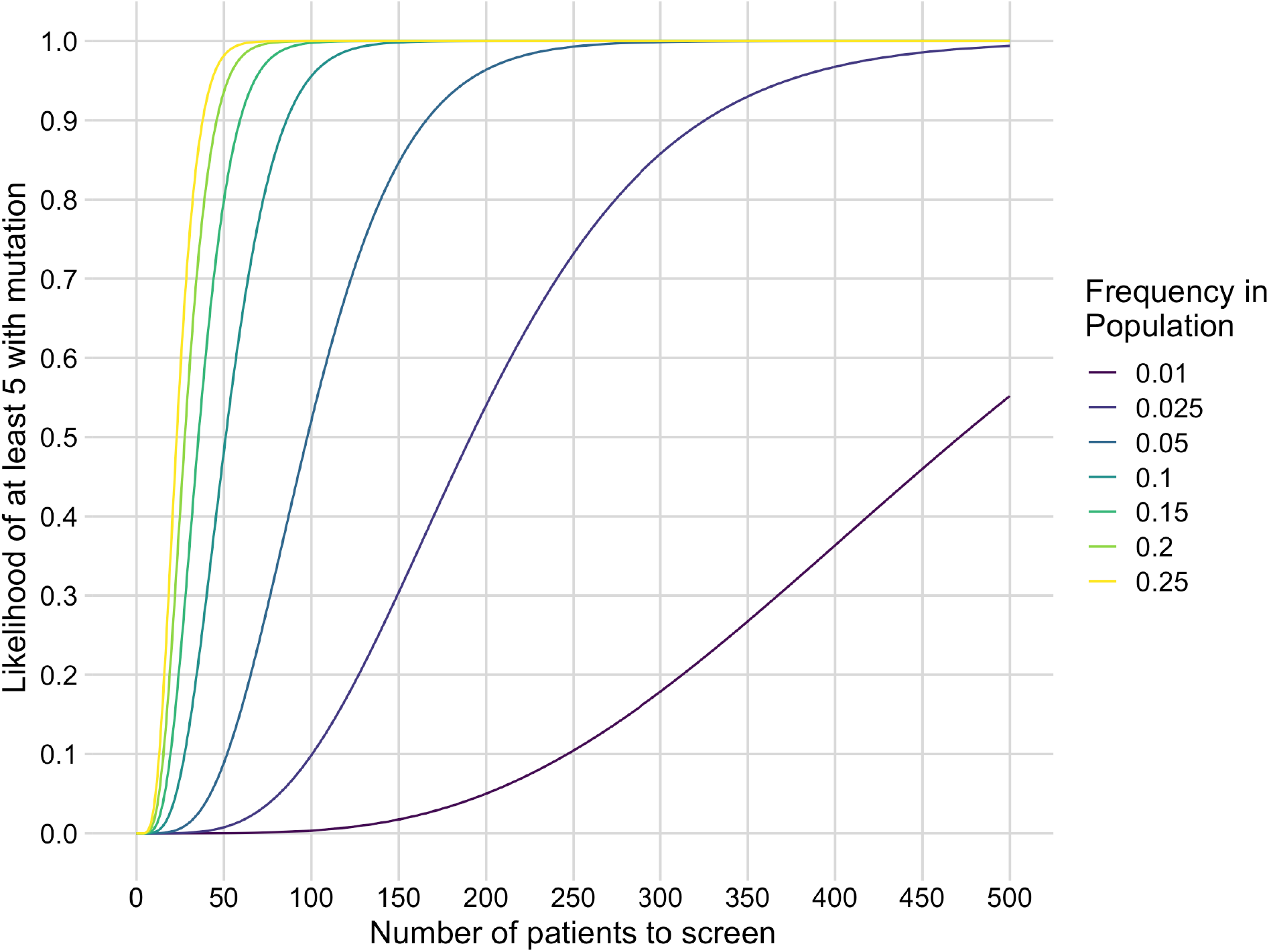
Power to identify at least 5 patients with a known driver mutation that is present in populations at varying low frequencies.

